# ezTrack: An open-source video analysis pipeline for the investigation of animal behavior

**DOI:** 10.1101/592592

**Authors:** Zachary T. Pennington, Zhe Dong, Regina Bowler, Yu Feng, Lauren M Vetere, Tristan Shuman, Denise J. Cai

## Abstract

Tracking small animal behavior by video is one of the most common tasks in the fields of neuroscience and psychology. Although commercial software exists for the execution of this task, commercial software often presents enormous cost to the researcher, and can also entail purchasing specific hardware setups that are not only expensive but lack adaptability. Moreover, the inaccessibility of the code underlying this software renders them inflexible. Alternatively, available open source options frequently require extensive model training and can be challenging for those inexperienced with programming. Here we present an open source and platform independent set of behavior analysis pipelines using interactive Python (iPython/Jupyter Notebook) that researchers with no prior programming experience can use. Two modules are described. One module can be used for the positional analysis of an individual animal across a session (i.e., location tracking), amenable to a wide range of behavioral tasks including conditioned place preference, water maze, light-dark box, open field, and elevated plus maze, to name but a few. A second module is described for the analysis of conditioned freezing behavior. For both modules, a range of interactive plots and visualizations are available to confirm that chosen parameters produce results that conform to the user’s approval. In addition, batch processing tools for the fast analysis of multiple videos is provided, and frame-by-frame output makes aligning the data with neural recording data simple. Lastly, options for cropping video frames to mitigate the influence of fiberoptic/electrophysiology cables, analyzing specified portions of time in a video, and defining regions of interest, can be implemented with ease.

## Introduction

The ability to process videos of small animals and automatically score their behavior is crucial to a number of common tasks in the fields of neuroscience and psychology – be it measuring the locomotor activity of an animal, defining its position in an arena, quantifying its interactions with an object, or assessing its engagement in defensive behaviors like freezing. Still, despite the nearly ubiquitous need for automated video analysis of this sort, there are substantial barriers to accessing these functions. One is cost – existing commercial software can cost several thousand dollars. Another is flexibility – commercial software often constrains the experimenter to particular hardware, operating systems, and video file types. The last is usability – while often powerful, existing free software can sometimes require substantial programming experience to implement and can involve complex algorithms (Sommer et al., 2011; Mathis et al., 2018).

To overcome these hurdles, we developed a simple, free, and open-source video analysis pipeline that 1) is accessible to those who have no programming background, 2) provides a wide array of interactive visualizations, 3) requires a minimal number of parameters to be set by the user, 4) produces tabular data in accessible file formats (e.g. csv), 5) accepts a large number of video file formats, and 6) is operating system and hardware independent. At the same time, being open-source, it allows users to modify the underlying code as they see fit.

Our behavior analysis pipeline, ezTrack, has two modules. The first is a module for analyzing an animal’s location throughout a session, the distance it travels, and the time it spends in user-defined regions of interest (ROI). The second allows the user to analyze freezing behavior, most relevant to the study of fear. For both modules, options for outputting frame-by-frame data as well as time-binned summary reports are provided. Additionally, both modules allow the user to either process individual videos with extensive visualizations of the results to aid in parameter selection, or to process large numbers of files simultaneously in a batch. Lastly, users can easily crop the frame of their videos and define the range of frames to be processed, in order to remove the influence of cables or fibers attached to the animal or other unwanted objects that might enter into the field of view. With this simple toolkit, a vast amount of automated behavioral analysis is readily performed.

## Results

ezTrack was designed to be implemented in iPython/Jupyter Notebook. Using Jupyter Notebook, the code is organized into “cells” – discrete, ordered sections that can be independently run by the user. Critically, instructions that inform the user what each cell does from a conceptual standpoint as they step through the code, as well as how to modify code when needed, precede each cell of code. That said, the core algorithms are implemented in separate python scripts (.py files), so that inexperienced programmers only have to set the values of a few key variables/parameters (e.g. the folder in which files are stored or a threshold value), and choose whether they want to run a particular cell or not. This makes the user interface balanced between usability and flexibility – the user can understand the algorithms conceptually without reading all of the code, while maintaining complete freedom to modify the algorithms if desired (**Supplementary Video 1**). Moreover, the output of running each cell are displayed directly below it so that the user can view the results of each cell of code that they run. To further leverage this, we provide numerous interactive plots and videos at critical steps to visually show the user the effect of algorithms/parameters, making the use of code intuitive. Tutorials of how to step through the code are presented in **Supplementary Video 1** (Location Tracking Module) and **Supplementary Video 2** (Freeze Analysis Module).

### Location Tracking Module

ezTrack’s Location Tracking Module assesses an animal’s location across the course of a single, continuous session. The Location Tracking Module can be used to calculate the amount of time the animal spends in user-defined ROIs, as well as the distance that it travels. It uses the animal’s center of mass to determine the animal’s location in each frame of the video (See **Supplementary Video 1** for tutorial and **Supplementary Video 3** for tracking example).

To validate that ezTrack’s Location Tracking Module works for a wide range of behavioral assays, we analyzed videos of mice being tested for their preference of a cocaine-paired chamber (conditioned place preference), their preference for the darker side of a two-chamber box (light-dark test), their preference for the closed arms of an elevated plus maze, and their preference for the quadrant of a water maze that formerly contained a hidden platform (Morris water maze) (Fig 1). As can be seen in Figure 1, ezTrack was able to track the position of animals in all of these assays, despite different lighting conditions, arena sizes, and camera orientations. It is particularly noteworthy that ezTrack worked well with the light-dark box because using commercial software, it is frequently difficult to maintain tracking of an animal when background lighting conditions change, or when there is generally low contrast between the foreground and background. Moreover, as demonstrated in **Supplementary Video 3**, ezTrack is quite robust to other objects that might enter the field of view. Provided the interfering object does not directly overlap with the animal in the field of view, tracking is maintained.

**Figure 1:**
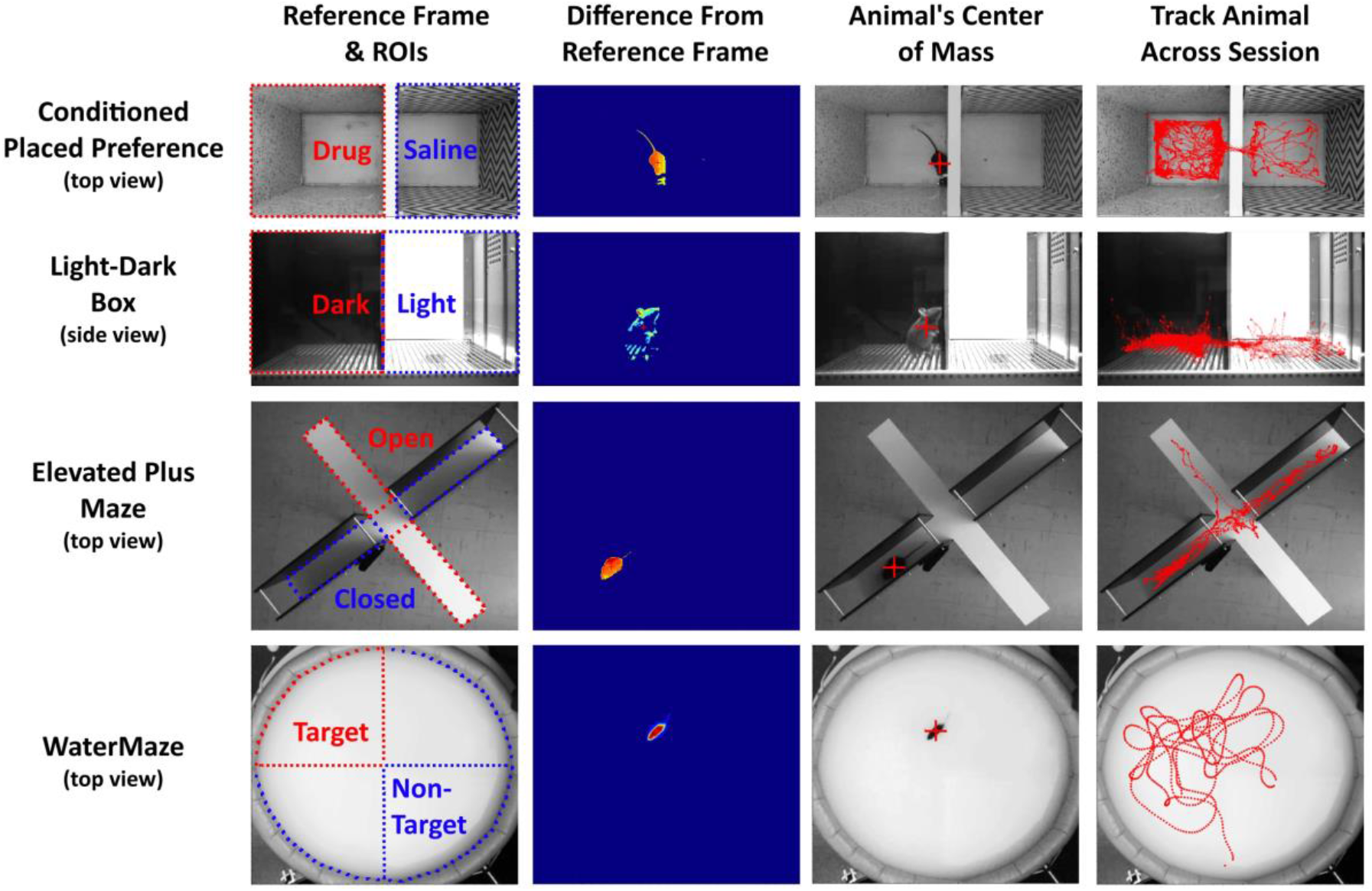
ezTrack Location Tracking Module. Using ezTrack’s Location Tracking Module, an animal’s center of mass is tracked across the session. After defining a reference frame and ROIs with a drawing tool (left-most panel), each frame is compared to the reference frame, taking their absolute difference (2nd panel). From these differences, the animal’s center of mass is calculated (3rd panel, crosshair indicates center of mass). The animal’s center of mass is then saved, as well as displayed atop the reference frame for visual inspection of the results (4th panel). All images come directly from ezTrack output.

To provide a quantitative validation of the Location Tracking Module, using ezTrack’s ROI drawing tool, we assessed the amount of time animals spent in ROIs for each task, and compared this with results of manual scoring. As can be seen in Figure 2A-F, there was a high correlation between automated and manual scoring for cocaine place preference (R^2^=0.99, y=1.2+1x), light-dark test (R^2^=97, y=6.9+0.94x) and elevated plus maze (R^2^=0.98, y=1.5+0.99x). Another useful tool provided by ezTrack’s Location Tracking Module is its calculation of the distance an animal moves on a frame-by-frame basis (in pixel units), derived by taking the Euclidean distance of the animal’s center of mass from one frame to the next. We examined conditioned place preference training data, in which animals were given either saline or cocaine. Using ezTrack, we were able to clearly track cocaine-induced hyperlocomotion (Fig 2G-H). Beyond many researchers’ interest in locomotion as an experimental variable, the frame-by-frame trace of distance that is automatically output by ezTrack (Fig 2H) is also useful in detecting anomalies in tracking. If something enters into the field of view and biases the center of mass, a large deflection in the distance traveled is anticipated. In this way, ezTrack alerts users to potential failures of tracking, a feature often not provided by other software.

**Figure 2:**
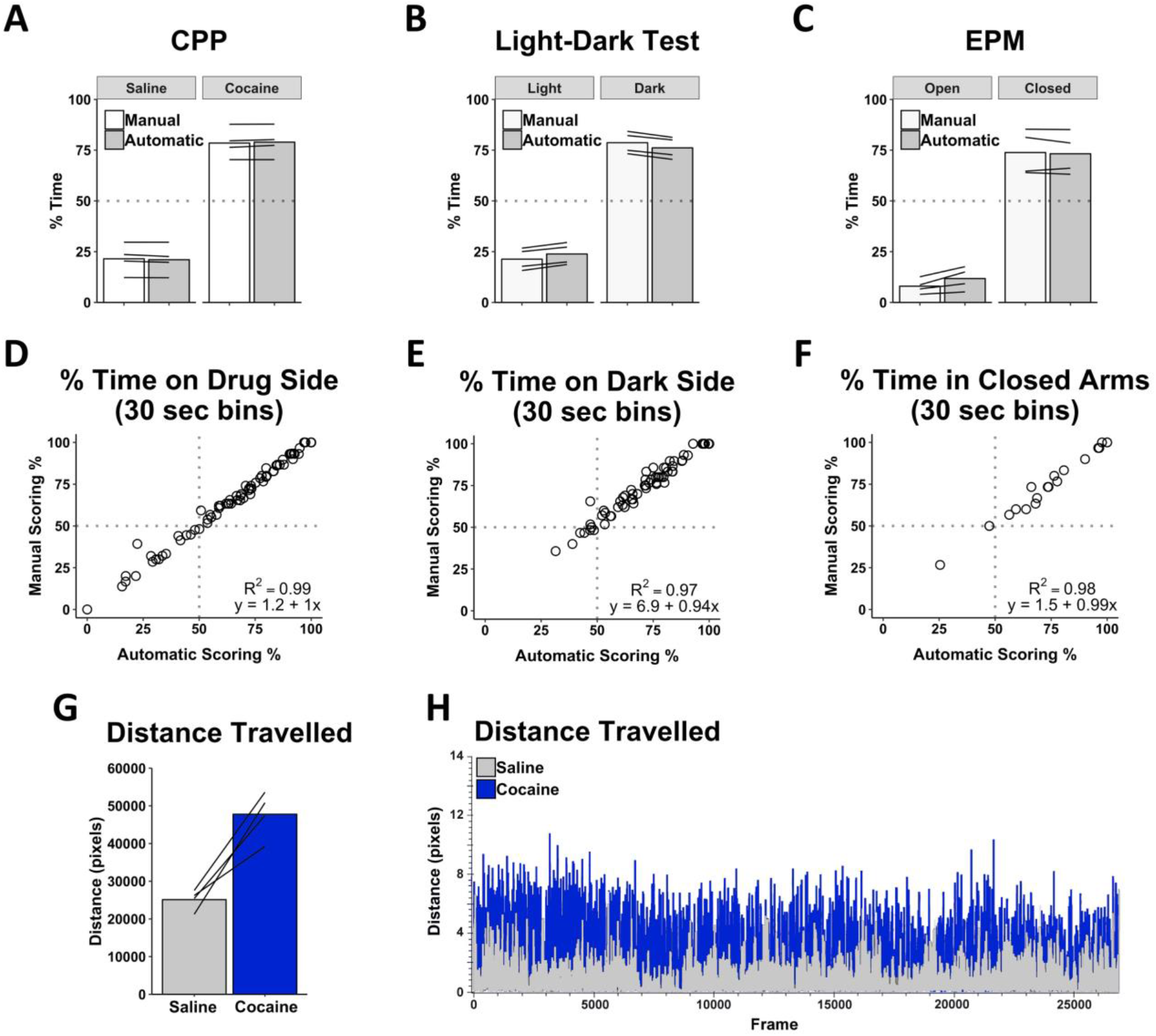
ezTrack Location Tracking Module Validation. Automated analysis of cocaine conditioned place preference (n=4 animals), light-dark box (n=4 animals), and elevated plus maze (n=4 animals) rendered nearly identical results as scoring by hand, both when looking at mean session data (A-C), and when session data was broken down into smaller time segments (E-F), demonstrating ezTrack’s accuracy. Lines in A-C represent individual subjects. G) Additionally, ezTrack reliably detects changes in locomotion, with increased motion in conditioned place preference training sessions when animals are given cocaine (each line represents an individual animal). H) Example plot from ezTrack showing frame by frame distance travelled during sessions when the same animal was given saline and cocaine injections. These plots are also useful to detect aberrations in tracking.

By default, ezTrack’s Location Tracking Module outputs frame-by-frame data in convenient csv files, making alignment with neurophysiological recordings a simple task. As a demonstration of this, we aligned single-photon *in vivo* calcium imaging with a Miniscope recording of hippocampal sub-region CA1 with location tracking results obtained from video of a mouse running back and forth on a 2meter linear track. As can be seen in Figure 3, We were able to define the location of each calcium event along the location of the linear track, and from this discern each cell’s place field (O’Keefe, 1976).

**Figure 3:**
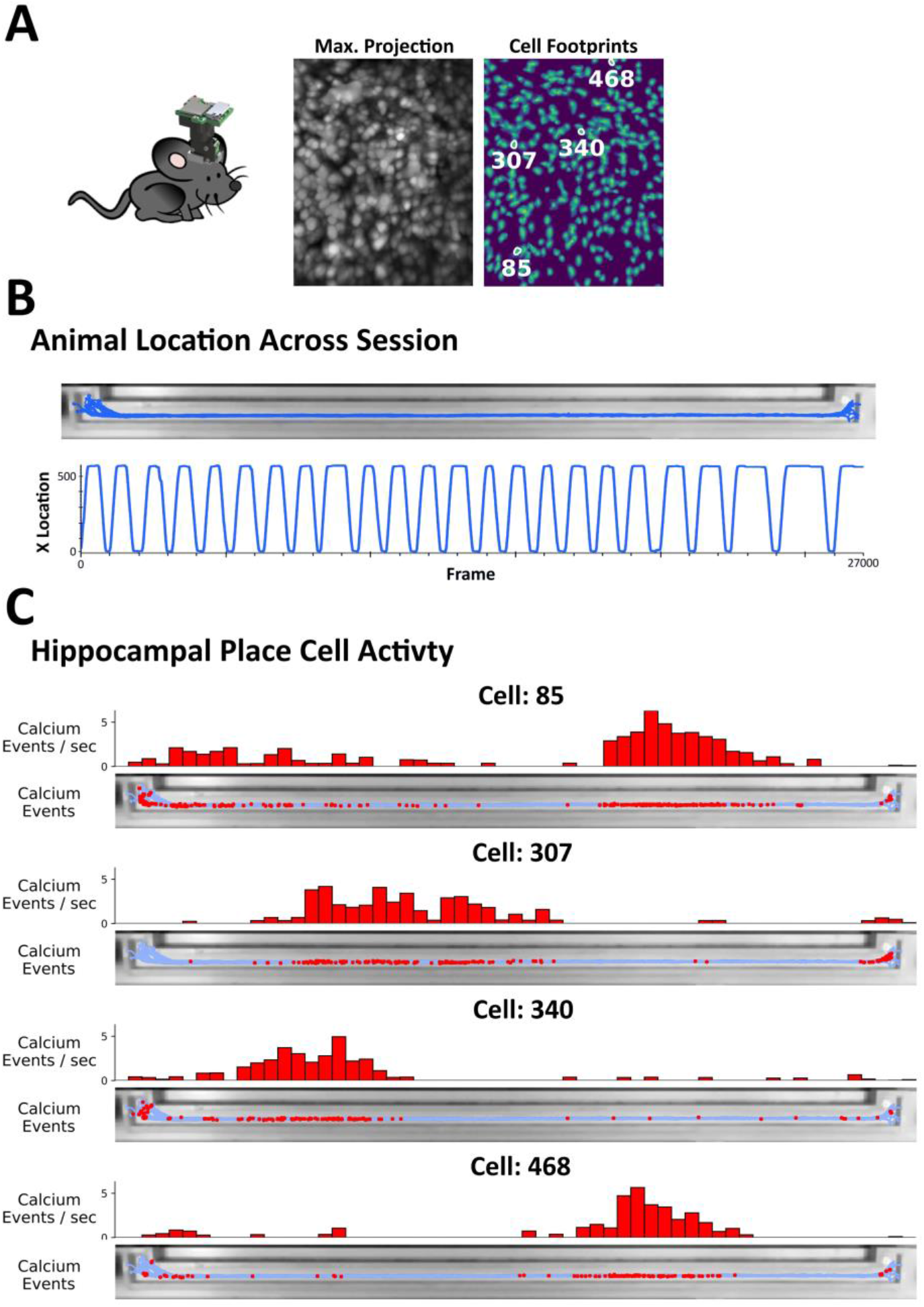
Alignment of Neurophysiological Data with ezTrack Results. A single mouse ran back and forth on a linear track wearing a Miniscope in order to track hippocampal place cells. A) Animal with head-mounted miniscope. Image on left shows max projection of Miniscope recording (for each pixel, max. value across session). Image on right shows isolated cell locations. Exemplary ‘place cells’ are highlighted. B) Using ezTrack, animal’s spatial position was examined during a single session. Top image shows animal position superimposed on linear track. Bottom image shows animal’s x position across a session. Note smooth tracking throughout. C) After deconvolving calcium activity, calcium events were aligned with spatial position to identify putative place cells. Top histogram shows normalized cell activity across track. Bottom image shows imposition of calcium events atop linear track.

### Freeze Analysis Module

ezTrack’s Freeze Analysis Module allows the user to measure freezing, employing its most common definition: the absence of movement, barring respiration. Similar to commercial automated software like VideoFreeze and FreezeFrame (Anagnostaras et al., 2010), ezTrack first measures the motion of the animal by assessing the number of pixels whose grayscale change exceeds a particular threshold from one frame to the next. Subsequently, an animal is scored as freezing when motion is below a user-defined threshold for a user-specified amount of time. The Freeze Analysis Module provides extensive visualizations in order to set thresholds which conform to manual inspection of videos. This includes interactive graphs in which the user can view the interrelationship of motion and freezing (Fig 4), as well as videos that allow the user to see what is being picked up as motion/freezing (**Supplementary Video 4**). Additionally, provided side-view recording is performed, ezTrack’s point and click cropping tool allows one to remove the influence of any fiberoptic/electrophysiology cables attached to the animal (Fig 4), which can continue to move even when the animal is freezing and bias measures of freezing. A tutorial of how to use the Freeze Analysis Module is provided in **Supplementary Video 2**.

**Figure 4:**
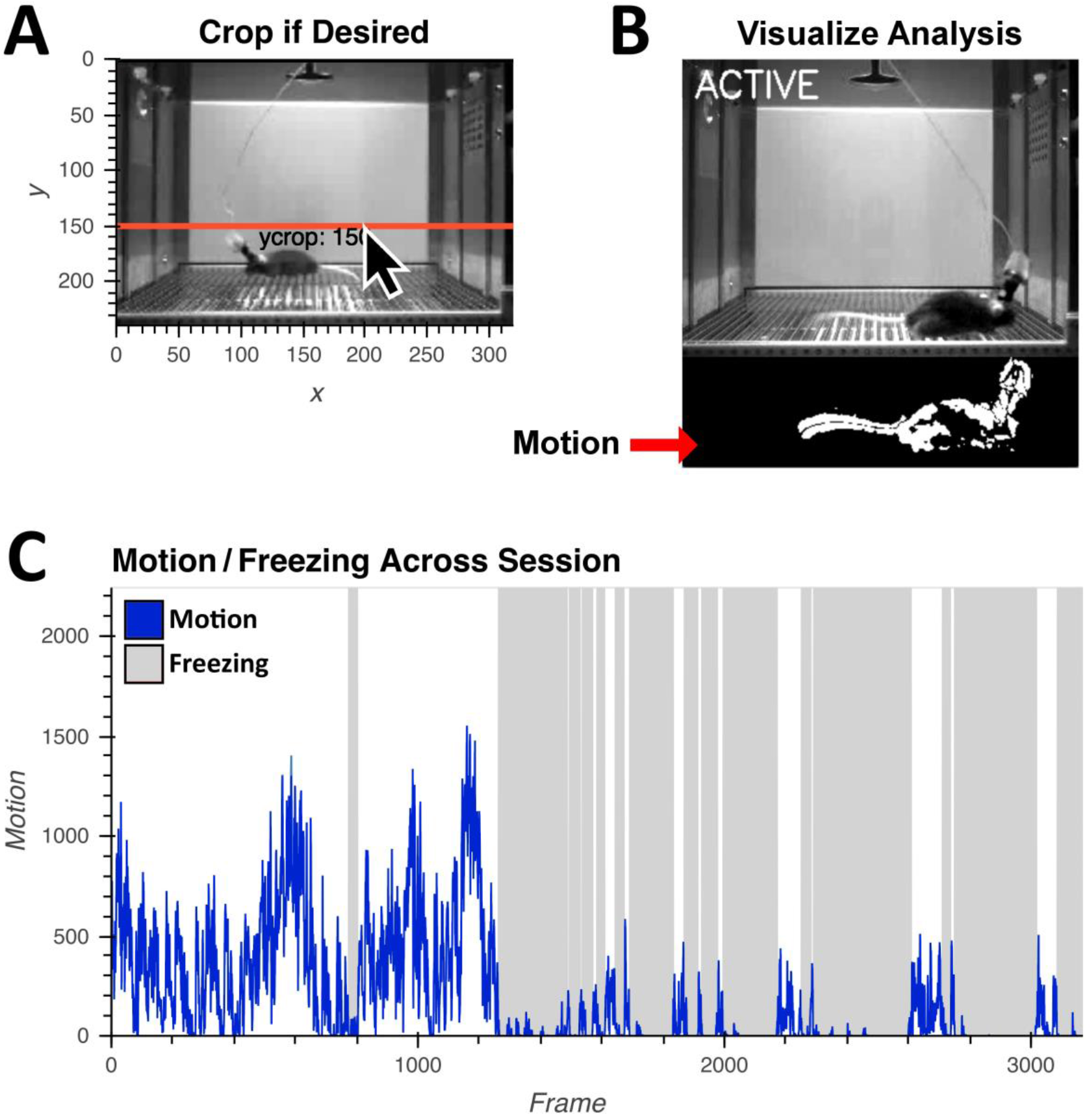
ezTrack Freeze Analysis Module. Using ezTrack’s Freeze Analysis Module, an animal’s motion and freezing are processed. A) Using point and click options, ezTrack’s Freeze Analysis Module allows the user to crop the top of the frame to remove the influence of cables. B) After analyzing data, segments can be played back to visualize scoring of motion and freezing (See Supplementary Video 4). Parameters – a motion threshold, freezing threshold, and minimum freeze duration – can be adjusted to conform to the experimenter’s judgment. C) Motion (blue) and freezing (gray) is plotted by ezTrack and frame-by-frame and binned summary data can then be saved to .csv files.

To validate accuracy of the Freeze Analysis Module, we analyzed videos from animals that underwent associative fear conditioning. Animals had undergone a varying number of conditioning trials (0, 1 or 3 footshocks; each 1 mA, 2 sec in duration) and were then placed back into the conditioning chamber the subsequent day to assess their freezing as an indicator of their fear memory. We compared the percentage of time spent freezing, determined either by manual scoring or using ezTrack, for the final test session. Ratings of average freezing for the entire 5-minute session were tightly correlated, as were ratings when we divided the test session into 30 second bins (Fig 5). Moreover, using both methods, ordinal increases in freezing paralleled the number of shocks received, and the relative magnitude of each animal’s freezing relative to the other animals was preserved (Fi 5). Of note, these animals were wearing head mounted Miniscopes (Cai et al., 2016) and had a cable extending from the top of the scope during recording. Many researchers have resorted to hand-scoring similar videos because commercial software is unable to remove the influence of cables. By taking the simple solution of cropping the top of the video, ezTrack allows the user to circumvent this problem.

**Figure 5:**
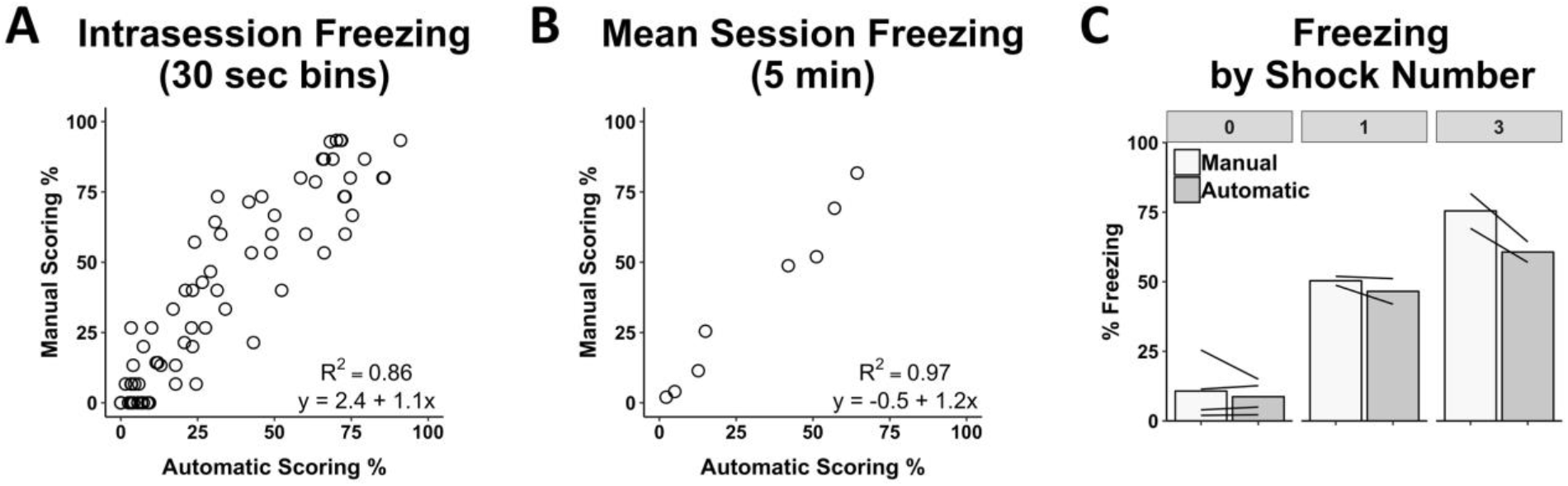
ezTrack Freeze Analysis Module Validation. The results of ezTrack’s Freeze Analysis Module were compared to those obtained from manual scoring of freezing. Automated analysis of freezing behavior was highly correlated with manual freezing, both when data was examined A) using 30 second time bins, B) and average freezing across the 5 min session (n=8 animals). C) Moreover, both manual and automated scoring show increased freezing with the number of shocks animals previously received, and relative between-mouse freezing levels are preserved (each mouse is represented by a line).

## Discussion

Above, we describe ezTrack, a flexible toolkit for the analysis of behavior, and demonstrate that ezTrack’s results highly correlate with manual scoring of an array of behavioral tasks, under varied experimental conditions. The tasks presented should be thought of as a small but representative sample of the vast number of behaviors that could be analyzed with ezTrack. The Location Tracking Module could be used for essentially any task in which an animal is moving throughout a stable background (e.g. place preference, water maze, radial arm maze, infinity maze, open field, etc). Moreover, it is conceivable that the Freeze Analysis Module could also be used to analyze other behaviors such as immobility in the forced swim task, or diurnal activity. Additionally, although ezTrack was tested here with mice, it could be used with any experimental species where the goal is to track the full position of the animal. In combination with the fact that ezTrack is compatible with a wide array of video types and operating systems, and requires no prior programming experience, we believe that ezTrack will be an easy and free tool to implement for many researchers.

ezTrack represents a substantial deviation from prior solutions to the problem of automated video analysis. The predominant solution – commercial software – has been relatively easy to use but costly and inflexible. Researchers are frequently bound to use particular hardware which may not always be compatible with their research questions. Moreover, what is done ‘under the hood’ is often proprietary and left unexplained. Alternatively, free solutions like DeepLabCut and ilastik (Sommer et al., 2011; Mathis et al., 2018) are great because they have the ability to track a wide range of sophisticated behaviors. However, they have been difficult to implement for scientists with little programming/computational experience. Moreover, although deep learning algorithms allow for the analysis of more complex data sets, they also require model training. ezTrack takes the middle road, providing a set of free tools that can be quickly implemented by anyone, allowing fast, automatic scoring of a seemingly endless number of data sets.

ezTrack will likely be particularly useful for researchers who combine behavioral monitoring with *in vivo* electrophysiological, optical and other biophysiological monitoring/manipulation techniques. First, commercial software does not readily output frame-by-frame data (e.g. Video Freeze), making it difficult to align biological and behavioral measurements at the necessary resolution. By default, ezTrack outputs this data in convenient csv files. Moreover, commercial software is frequently biased by cables that are attached to the animal. ezTrack permits the user to crop the frame of the video to remove the influence of these objects. Simple as they are, these features of ezTrack will make the research process much easier.

ezTrack is simple not only in its technical implementation, but in its conceptual nature (see Methods). Both the Location Tracking Module and the Freeze Analysis Module have very few parameters that must be set by the user and there are no hidden parameters in the underlying code. In combination with the rich set of visual outputs that ezTrack provides, it is easy for the researcher to understand the relationship between the data input and the data output, and thus, how parameters can be changed to improve tracking accuracy. The intuitive nature of ezTrack will provide transparency in its results in comparison to programs where data is processed within a ‘black box’. Furthermore, ezTrack’s simplicity facilitates modifications to the code for those who wish to do so. Along these same lines, ezTrack could be an incredible instructional tool for teaching students at many levels how to process video content.

Data reproducibility is another benefit of ezTrack. ezTrack’s use of Jupyter Notebook, an html-based platform, makes it so that all selected parameters, the steps that were taken to process a dataset, and the output, can be saved in a highly readable pdf or html file (being a printout of the notebook the data was processed in). This stands in contrast to graphical-user-interface-based systems in which point and click options are not necessarily saved, and when they are, are not convenient to access. This makes it easier for the experimenter and others to go back and confirm how the data was processed.

ezTrack is not without limitations. In its current form, ezTrack measures the activity of a single animal and works predominantly by tracking the animal’s center of mass. This occludes the analysis of social interaction, of more specific sorts of interactions with objects (e.g. sniffing, grabbing, burying), and of fine motor movements. It is likely that the measurement of some of these behaviors – particularly fine motor movements – are better accomplished with more sophisticated algorithms (Mathis et al., 2018). However, other features, like tracking an animal’s direction and tracking multiple animals, can be added to ezTrack in the future as needed. Indeed, a major benefit of ezTrack being open-source is that it is relatively fast to update and distribute new features and modifications. For example, due to the simplicity of the computation, ezTrack could easily be adapted to work in real-time in conjunction with output devices; researchers could adapt ezTrack so that when animals are in a particular location an output is triggered. Although we have not included this option in this version, these and additional modifications could be easily incorporated in future versions of this behavioral analysis pipeline. In closing, ezTrack can fulfill a broad set of researchers’ needs and the open source platform facilitates the flexibility of adding new functions in ezTrack.

## Methods

### Installation and Dependencies

Detailed instructions on ezTrack installation can be found on Github: github.com/denisecailab/ezTrack. In brief, after following the instructions for downloading Conda/Miniconda (a free package management system) the commands are given for installing both Jupyter Notebook and other free packages required for the code to be run. Subsequently, Python/notebook files can be downloaded from our github account and run within Jupyter Notebook. **Supplementary Videos 1 and 2** provide a brief tutorial for using Jupyter Notebook. The following opensource packages are used for video import, data manipulation, and interactive plotting: iPython (Perez and Granger, 2007), OpenCV (Bradski, 2000), Pandas (McKinney, 2010), HoloViews (PyViz, 2018), SciPy (Jones et al., 2001), Matplotlib (Hunter, 2007), Bokeh (Bokeh, 2018), and NumPy (Oliphant, 2006).

### Video Acquisition Suggestions

Although ezTrack works on a wide variety of videos, there are certain conditions that will render tracking unreliable. Acquired videos must be captured with a mounted camera so that the field of view is fixed throughout the recording session. Moreover, ezTrack works best under conditions where there is good contrast between the animal and the background – if it is hard for the human experimenter to see the animal, it will likely be problematic for ezTrack to calculate the center of mass of the animal and track its location. ezTrack was programmed to be robust against the influence of objects other than the animal in the field of view – cropping of the video frame is supported, and the Location Tracking Module has a function that reduces the influence of objects that might enter the field of view by weighting the region surrounding where the animal was on the prior frame (described in detail below). Nevertheless, ezTrack will work best when only the animal and a stable environment are in the field of view. Moreover, the animal should not be able to exit and re-enter the field of view. Also, in its current state ezTrack only tracks a single animal. Simultaneously recording multiple arenas with one camera is possible; however, for now, the experimenter will have to individually analyze each arena by cropping the other arenas out of the field of view. All batch processing features assume that the position of the camera, the position of the environment within the field of view, and the lighting conditions, are consistent across sessions. We have tested ezTrack with wmv, avi (mp4 codec), and mpg1 videos, but many more video filetypes are possible. If your particular filetype is not supported, there are plenty of free video converters available online. Lastly, note that high definition videos can take longer to process and down sampling/compression of video may be desired.

### Location Tracking Module

#### General Description

The Location Tracking Module tracks the center of mass of an animal within the field of view on a frame-by-frame basis. From this, the distance the animal travels and its time spent in particular ROIs, are calculated. Video can be taken either looking down on the animal, or from the side, though side view recording can put distance measurements on an arbitrary scale. Using a drawing toolbox, the user is able to crop the field of view, and the user can also restrict the analysis of the video to a particular range (e.g. frames 500-1000). Additionally, the user is able to draw an unlimited number of ROIs, and for each frame ezTrack will determine if the animal is in a particular region or not. Notably, regions of interest are allowed to overlap, allowing sub-regions to be analyzed. While running the code, interactive plots are generated which allow the user to view both the distance travelled across the session and traces of where the animal went. As an added feature, if video files are small (e.g. 100 × 100 pixels), or if they are odd dimensions (e.g. 30 × 1000 pixels), ezTrack allows the user to stretch the video horizontally/vertically for presentation purposes. The frame by frame location of the animal in x, y coordinates, as well as whether the animal is in each defined region of interest, are saved to a csv file. If the user wants summary information, options are provided for specifying time bins, and a summary file will be generated giving the distance travelled in each time bin, as well as the proportion of time spent in each defined ROI. Once the user is comfortable that tracking is working well for individual videos, they can then perform batch processing, in which every video file in a folder will be processed.

#### Methods for Location Tracking

Location tracking is accomplished by comparing each frame in a video to a reference frame where the animal is not present in the environment. ezTrack supports two ways to define this reference frame. The easier, faster and often more reliable option is to generate a reference frame from the video being analyzed (i.e. the video with the animal). ezTrack will generate a reference frame based upon a random sample of frames across the video (default = 100). For each pixel in the field of view, the median value across the selection of frames is taken. Critically, this median image will not contain the animal unless the animal is in one location for over 50% of the session. If this occurs, the user can alternatively supply a separate video of the same environment with the same lighting conditions and with the environment in the same position in the field of view without the animal. When batch processing, the first option is always used, such that the reference frame for each video is generated from that video

Next, to determine the center of mass for the animal, for each frame, the absolute grayscale intensity difference from the reference frame is calculated on a pixel-by-pixel basis. In order to mitigate the influence of random, low intensity fluctuations in pixel intensity values when calculating the center of mass, which we find can greatly bias accuracy, pixel differences are thresholded so that only the difference values above a user-defined percentile of all pixel differences are considered. We have used the 99^th^ percentile and this has worked incredibly well. Notably, because this criterion scales relatively, the animal will generally be tracked equally well if it moves from a high to low contrast area of the arena, as in the light-dark box (see Fig 1). Lastly, the center of mass on these values is calculated, returning the x,y coordinates of the animal.

What happens if something enters the field of view, biasing the center of mass to be between the animal and the object in question? ezTrack allows the user to weight the pixels surrounding the animal’s previous location to avoid this. For each frame, a weight (w) can be applied to a square window surrounding the center of mass of the animal on the previous frame, the size of which is set by the user. Pixel difference values outside of the window are multiplied by (1-w), where w is on a scale from 0-1. Thus, if the w is set by the user to 1, pixel value differences that are outside the window will be set to zero. By contrast, if w is set to 0, the interior/exterior of the window carry equal weight. In this way, the influence of pixel changes outside the window can be restrained to a minimal level controlled by a single parameter, w. To mitigate the potential issue that the animal moves out of the window (such as when the recording starts before the animal is placed in the field of view), w can also be set to something between 0-1, which will allow the window to ‘snap’ back to the animal. We have been using 0.8-0.9 with good results.

Lastly, the distance the animal traveled from one frame to the next is calculated as the Euclidean distance between the center of mass on adjacent frames.

Notably, the only parameter required to be set by the user is the percentile for setting pixel difference values to zero. The other two parameters – setting a weight and a window size for weighting difference values in the vicinity of the animal’s location on the prior frame – are optional.

### Freeze Analysis Module

#### General Description

In order to detect freezing, ezTrack first measures the motion of an animal by calculating the number of pixels whose grayscale value changes from one frame to the next. However, because most videos display many small fluctuations in pixel values, even with a static scene, a cutoff must first be set to separate pixel changes that are attributable to the animal moving versus those that would occur with no animal in the box. ezTrack provides a calibration tool for examining the distribution of grayscale changes for a video of an empty box and provides a suggested cutoff. The user can also restrict the analysis of the video to a particular range of frames. After measuring the motion of an animal, freezing can be calculated by assessing the amount of time the animal’s motion drops below a user-defined threshold. See **Supplementary Video 2** for a tutorial.

#### Methods for Freeze Analysis

##### Calibration

To alleviate small fluctuations in pixel values attributable to noise, we found it helpful to first implement a gaussian filter (sigma=1) on each image. Then, we calculate pixel-wise differences on consecutive frames, and count the number of pixels whose difference value exceed a certain cutoff (the motion threshold, or MT). To define MT, the user provides a short video of the recording environment without an animal (~10 sec). ezTrack will then calculate the distribution of pixel grayscale changes across this time period. It is then possible to set MT based on this null distribution. With our particular setup, we found that twice the 99.99^th^ percentile worked well. However, ezTrack provides a histogram of the distribution of difference values and also provides the user with the ability to see what is being picked up as motion side by side with the original video for a given MT (**Supplementary Video 4**). If the user senses too much noise or not enough signal is being detected, they can modify MT as they see fit. The only recommendation is that they maintain this threshold across all animals in an individual experiment. Provided video settings are not changed across days, we find that the MT for a given environment is very stable.

##### Measuring Motion/Freezing

Once the user defines MT, they can then analyze motion and freezing on an individual session. Each frame is gaussian filtered and the number of pixels whose grayscale change relative to the prior frame exceeds MT is determined. The user can then set a threshold for the number of changed pixels below which freezing is declared – the freezing threshold, or FT. As an additional criterion, a minimum freeze duration can be set (e.g. half a second). Thus, for any given succession of frames in which the number of changed pixels falls below FT for a period of at least the minimum freeze duration, the animal will be marked as freezing. The freezing threshold can be set by the user by first inspecting plots of motion (i.e. changed pixels) across the session and noting the values corresponding to markedly low points, presumably freezing. After selecting a tentative FT, they can then watch video in which they can inspect the accuracy of their threshold and adjust it as they see fit (**Supplementary Videos 2 and 4**). When the user is satisfied with a set of parameters, they can then proceed to process several videos in batch. When batch processing, individual csv files containing frame-by-frame motion and freezing values are saved for each video. Additionally, the user can define time bins and a summary file containing the average freezing and motion during each time bin, for each video, will be returned.

### Behavioral Testing and Manual Scoring

#### Manual Scoring

All manual scoring was performed using a time-sampling procedure in which the instantaneous location/freezing of an animal was assessed every 1-2 sec (2 sec for freezing, 1 sec for all other behaviors). Categorical responses were then turned into proportions by examining several responses over time.

#### Animals

Adult male C57/BL6J mice from Jackson Laboratories were used for all testing. All animal procedures were approved by Mount Sinai’s IACUC.

#### Behavior

##### Fear Conditioning

Animals were placed in a conditioning chamber with a distinct scent and received either 0, 1, or 3 footshocks. Subsequently, animals were placed back into this environment for a 5 minute test session in which no shocks were given. Automated scoring with ezTrack used a 0.5 second minimum freeze duration and the tops of videos frames were cropped to remove the influence of Miniscope cables. The amount of time spent freezing was assessed. The same freezing threshold was used for all animals, which was determined by visual inspection of a couple of animals.

##### Conditioned Place Preference

Conditioning took place across two days. On each day, animals were confined to one side of a conditioning chamber for 15 minutes immediately after receiving an injection of saline or cocaine (15 mg/kg, i.p.). They were placed on alternate sides across training sessions so that one side would be associated with cocaine. The following day, animals were placed back in the chamber for 15 minutes and allowed to freely explore its two sides. The amount of time spent in the cocaine-paired side was examined. Additionally, distance travelled during training was measured.

##### Light-Dark Test

Animals were placed in a chamber with a small opening connecting its two sides, one side being brightly lit (~400 lumens), the other being dimly lit (~10 lumens). The front of the dark side, through which the animal was recorded, was covered in a red transparent film to prevent the spread of white light into the box; the front of the brightly lit side was translucent. In order to facilitate visibility of the animal, an infrared light and camera were used (hence, lighting in Fig 1 does not reflect perceived lighting of the animal). The proportion of time spent on the dark side was measured.

##### Elevated Plus Maze

Animals were placed at the center of an elevated, plus-shaped, platform with two walled arms (closed) and two unwalled arms (open) for two and half minutes. The proportion of time animals spent in the closed arms was assessed.

##### Water Maze

A circular tub (4ft in diameter) filled with a combination of water and white paint. A platform (10cm in diameter) was submerged in one quadrant and start locations were randomized to one of four equidistant locations. Animals were trained three times per day for six days. In each trial, mice were given 60 seconds to find the platform. If mice found the platform earlier than 60 seconds, the trial ended then. If mice failed to find the platform, the trial terminated at 60 seconds. After each trial, mice were put on the platform for 15 seconds.

##### Linear Track

For linear track recording sessions, a water-restricted animal ran back and forth on a 2-meter-long linear track during a 15 minute session in order to earn water rewards that were alternately given at each end. The data presented come from a single session for a well-trained mouse.

### Miniscope Imaging

A Miniscope was used to image calcium events in the CA1 region of the hippocampus as previously described (Cai et al., 2016; Shuman et al., 2018). The analysis of calcium recordings was performed using an in-house written script based on existing algorithms (Pnevmatikakis et al., 2016; Lu et al., 2018). The script used in this paper is available online: https://github.com/DeniseCaiLab/minian. The pipeline is described briefly here: First, the raw calcium imaging videos are passed through a median filter and subsequently an anisotropic filter frame-wise to reduce granular noise. A general background is estimated by convolving each frame with a mean kernel whose size is a few times larger than the expected size of cells, and this background is then subtracted from each frame to remove the vignetting effect. Next, a standard fft-based cross-correlation method is used to estimate and correct for translational motion in the video, followed by a demon-based algorithm to correct for residual non-rigid motion. Next, an over-complete set of “seed” pixels is generated by finding the local-maxima on the maximum projection of a randomly selected subset of frames. These seeds are further refined and merged based on properties of their corresponding temporal dynamics (e.g. peak-to-peak values or peak-to-noise ratios), after which these seeds are used to identify the initial regions of interest that may correspond to cells (Lu et al., 2018). Lastly, the seeds-based initial regions of interest and their corresponding activities are fed into the CNMF algorithm as initial spatial and temporal terms respectively (Pnevmatikakis et al., 2016). The CNMF algorithm decomposes the video into a matrix of spatial footprints and a matrix of corresponding temporal activities for each cell. In addition, the CNMF algorithm models the temporal activities of cells as an auto-regressive process, and de-convolves the signals to give an estimation of underlying spiking activity. Due to the slow evolution of calcium signals, we use the de-convolved signals for further analysis of place cells. Due to imperfect regularization of spike counts during the estimation of the deconvolved signal, there are many low-amplitude, false-positive spikes. Thus, we performed a thresholding on the signals to get a binary variable representing whether a calcium event was observed on given frames. This threshold was chosen arbitrarily by observing the distribution of the de-convolved signals across cells. To find cells that are most likely coding for the location of the animal, we align the behavioral tracking results to the calcium recordings frame-wise using a nearest-neighbor approach. Then, we calculate spatial information content for each cell by treating the thresholded, deconvolved signals as “spike trains” (Skaggs et al., 1992). Example cells with high spatial information content are shown in Figure 3.

## Notes

**Funding**: This work was supported by the McKnight Memory and Cognitive Disorders Award to DJC, the Klingenstein-Simons Fellowship to DJC, the Brain Research Foundation Award to DJC, NARSAD Young Investigator Award to DJC, the Botanical Center Pilot Award to DJC, the CURE Award to TS, the American Epilepsy Society Award to TS, the Friedman Scholar Award to DJC, NIDA 5T32DA007135 to ZTP, and 5T32AG049688-02 to LMF.

#### Summary of Updates

An additional figure demonstrating ezTrack's compatibility with physiological recordings has been supplied. Additionally, multiple supplemental tutorial videos have been added.

https://github.com/DeniseCaiLab

